# *Fusobacterium* genomics using MinION and Illumina sequencing enables genome completion and correction

**DOI:** 10.1101/305573

**Authors:** S. Michelle Todd, Robert E. Settlage, Kevin K. Lahmers, Daniel J. Slade

## Abstract

Understanding the virulence mechanisms of human pathogens from the genus *Fusobacterium* has been hindered by a lack of properly assembled and annotated genomes. Here we report the first complete genomes for seven *Fusobacterium* strains, as well as resequencing of the reference strain *F. nucleatum* subsp. *nucleatum* ATCC 25586 (seven total species, eight total genomes). A highly efficient and cost-effective sequencing pipeline was achieved using sample multiplexing for short-read Illumina (150 bp) and long-read Oxford Nanopore MinION (>80 kbp) platforms, coupled with genome assembly using the open-source software Unicycler. When compared to currently available draft assemblies (previously 24-67 contigs), these genomes are highly accurate and consist of only one complete chromosome. We present the complete genome sequence of *F. nucleatum* 23726, a genetically tractable and biomedically important strain, and in addition, reveal that the previous *F. nucleatum 25586* genome assembly contains a 452 kb genomic inversion that has been corrected using our sequencing and assembly pipeline. To enable the scientific community, we concurrently use these genomes to launch FusoPortal, a repository of interactive and downloadable genomic data, genome maps, gene annotations, and protein functional analysis and classification. In summary, this study provides detailed methods for accurately sequencing, assembling, and annotating *Fusobacterium* genomes, which will enhance efforts to properly identify virulence proteins that may contribute to a repertoire of diseases including periodontitis, pre-term birth, and colorectal cancer.

## Background & Summary

Multiple *Fusobacterium* species are oral pathogens that can infect a broad range of human organ and tissue niches^1,2^. *Fusobacterium nucleatum* has recently been connected with colorectal cancer (CRC)^3,4^, with studies showing this bacterium induces a pro-inflammatory tumor microenvironment^5,6^ and chemoresistance against drugs used to treat CRC^7^. Despite the importance of this bacterium in human diseases, there is a lack of completed genomes of biomedically relevant isolates to allow for virulence factor identification. Further motivation for complete sequencing and assembly of a library of *Fusobacterium* genomes came from the observation that our bioinformatic analysis frequently uncovered a high percentage of large, predicted secreted proteins (~3,000-11,000 bp) in the *F. nucleatum* 23726 genome that were missing critical protein domains at either the N- or C-terminus (e.g. N-terminal Sec signal sequences).

The genome of *F. nucleatum subsp. nucleatum* ATCC 25586, which is the standard *F. nucleatum* reference strain, was completed in 2002 using cosmid and *λ* phage technologies to achieve long reads (10-35 kb) and facilitate genome assembly^8^. More recently, several *Fusobacterium* draft genomes have been sequenced using short-read technologies (454 Life Sciences), presumably making complete genome assembly difficult due to repeat regions (e.g. CRISPR arrays, transposons). Similarly, Illumina sequencing is highly accurate and widely available, but is not optimal for assembling whole genomes because of read length limitations (~ 150 bp). With the emergence of next generation long-read sequencing (Pacific Biosciences, Oxford Nanopore Technologies MinION), assembling whole genomes is now becoming standard and affordable for academic research settings. The recent combination of MinION long-read and Illumina short-read technologies to scaffold and polish DNA sequencing data, respectively, has created a robust pipeline for microbial genome completion and subsequent gene identification and characterization^9^. A follow up study by these scientists detailed their methods for concurrently sequencing twelve *Klebsiella* genomes through multiplex sampling^10^. Following this experimental road map, we outline our experimental methods for the first completely sequenced, assembled, and annotated *Fusobacterium* genomes using MinION technology. In addition, these inaugural genomes are used to launch the FusoPortal genome and bioinformatic analysis repository. In summary, this study provides key resources to further determine how multiple *Fusobacterium* species contribute to a variety of human infections and diseases.

## Methods

### Bacterial growth and genomic DNA preparation

All strains of *Fusobacterium* were grown overnight in CBHK (Columbia Broth, hemin (5 *μ*g/ml), and menadione (0.5 *μ*g/ml) at 37 °C in an anaerobic chamber (90% N_2_, 5% CO_2_, 5% H_2_). Genomic DNA from stationary phase bacteria was isolated in diH_2_O from each strain using a Wizard isolation kit (Promega), and quantitated using a Qubit fluorimeter (Life Technologies).

### Short-read Illumina sequencing

Short-read DNA sequencing was carried out at the Genomic Sequence Center at the Virginia Tech Biocomplexity Institute and Novogene (strain *F. nucleatum* 25586). For sequencing at Virginia Tech, DNA-seq libraries were constructed using PrepX ILM 32i DNA Library Reagent Kit on an Apollo 324 NGS library prep system. Briefly 150 ng of genomic DNA was fragmented using a Covaris M220 Focused-ultrasonicator to 400 bp. The ends were repaired and an ‘A’ base added to the 3’ end for ligation to the adapters which have a single ‘T’ base overhang at their 3’ end. Following ligation, the libraries were amplified by 7 cycles of PCR and barcoded. The library generated was validated by Agilent TapeStation and quantitated using Quant-iT dsDNA HS Kit (Invitrogen) and qPCR. The libraries were then pooled and sequenced using a NextSeq 500/550 Mid Output kit V2 (300 cycles) (P/N FC-404-2003) to 2 × 150 cycles. BCL files were generated using Illumina NextSeq Control Software v2.1.0.32 with Real Time Analysis RTA v2.4.11.0. BCL files were converted to FASTQ files, adapters trimmed and demultiplexed using bcl2fastq Conversion Software v2.20. Illumina sequencing statistics and genome coverage are detailed in Table 1, and the public availability of the data at NCBI is detailed in Table 4.

**Table 1.** Statistics for short read Illumina sequencing.

| Species | Strain | # Illumina Sequences | Base Pairs | Mean Length | Max Length | Genome Size | Mean Depth |
| --- | --- | --- | --- | --- | --- | --- | --- |
| *F. nucleatum* | 23726 | 5,163,442 | 774.5 mb | 150.2 | 151 | 2,299,539 | 336 X |
| *F. nucleatum* | 25586 | 1,250,000 | 187.5 mb | 150.0 | 150 | 2,180,101 | 86.0 X |
| *F. varium* | 27725 | 1,250,000 | 187.5 mb | 150.2 | 151 | 3,303,644<br>Plasmid:<br>42,814 | 56.8 X |
| *F. ulcerans* | 49185 | 1,250,000 | 187.5 mb | 150.2 | 151 | 3,537,675 | 53.1 X |
| *F. mortiferum* | 9817 | 1,250,000 | 187.5 mb | 150.2 | 151 | 2,716,766 | 69.2 X |
| *F. gonidiaformans* | 25563 | 1,250,000 | 187.5 mb | 150.2 | 151 | 1,678,881 | 111 X |
| *F. periodonticum* | 2_1_31 | 1,250,000 | 187.5 mb | 150.2 | 151 | 2,541,084 | 73.8 X |
| *F. necrophorum* | 1_1_36S | 1,250,000 | 187.5 mb | 150.2 | 151 | 2,286,018 | 82.2 X |

### Long-read MinION sequencing

Purified *Fusobacterium* genomic DNA was sequenced on a MinION sequencing device (Oxford Nanopore Technologies) using the 1D Genomic DNA sequencing kit SQK-LSK108 according to Oxford Nanopore Technologies instructions. Multiplexed samples were barcoded using the 1D Native Barcoding Kit (EXP-NBD103) according to instructions. Briefly, purified genomic DNA was repaired with NEBNext FFPE Repair Mix (New England Biolabs). The NEBNext Ultra II End-Repair/dA-tailing Module was utilized to phosphorylate 5’ ends and add a deoxyadenosine monophosphate (dAMP) to the 3’ ends of the repaired DNA. For multiplexed samples, barcodes were ligated to the end-prepped DNA using the NEB Blunt/TA Master Mix (New England Biolabs). Barcoded samples were pooled into a single reaction and an adapter (Oxford Nanopore Technologies) was ligated to the DNA using the NEBNext Quick T4 DNA Ligase (New England Biolabs). For single reactions, an adapter (Oxford Nanopore Technologies) was ligated to the end-prepped DNA using the NEB Blunt/TA Master Mix (New England Biolabs). The DNA was purified with AMPureXP beads (Beckman Coulter, Danvers, MA) following each enzymatic reaction. Purified, adapted DNA was sequenced on a MK1B (MIN-101B) MinION with a FLO-MIN106 (SpotON) R9.4 or FLO-MIN107 (SpotON) 9.5 flow cell using MinKNOW software version 1.7.10 or 1.7.14 (Oxford Nanopore Technologies). After sequencing, Fast5 files were basecalled using Albacore version 2.1.7 (Oxford Nanopore) on a Macbook Pro with a 3.3 GHz Intel Core i7 processor. For multiplexed samples, basecalled fastq files were demultiplexed based on the ligated barcode using Porechop and adaptors were trimmed. Sample preparation and sequencing details are described in Table 2, and the MinION sequencing statistics and genome coverage are detailed in Table 3. As an example of data quality, Figure 1 shows the long read coverage obtained using MinION sequences for the *F. necrophorum funduliforme* 1_1_36S genome.

**Figure 1:**
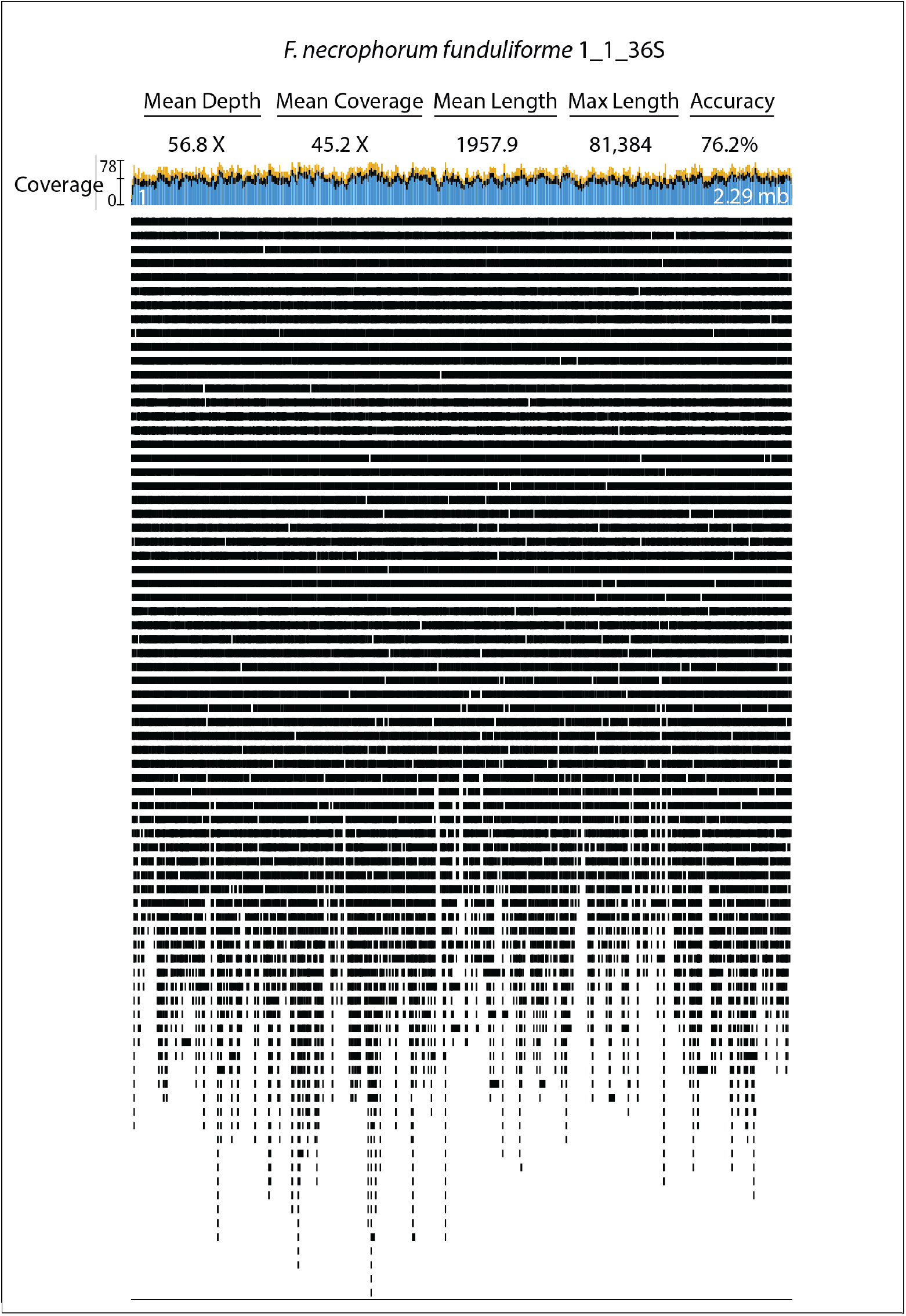
Statistics and Mapping of *F. necrophorum funduliforme* 1_1_36S MinION long-reads. Post complete genome assembly, MinION reads were mapped to the *F. necrophorum funduliforme* 1_1_36S genome using Geneious version 9.1.4 software.

**Table 2.** Experimental details for MinION sequencing.

| Sample IDs | Sequencing Kit | Flow Cell | MinKNOW Version | Input DNA (μg) |
| --- | --- | --- | --- | --- |
| 23726 | SQK-LSK108 | FLO-MIN106 (SpotON) R9.4 | 1.7.10 | 2.38 |
| 25586 | SQK-LSK108 | FLO-MIN107 (SpotON) R9.5 | 1.7.14 | 3.96 |
| 49185, 9817, 25563, 2_1_31, 1_1_36S, 27725 # | SQK-LSK108, EXP-NBD103 | FLO-MIN107 (SpotON) R9.5 | 1.7.14 | 3.72, 3.12, 2.94, 0.6, 2.31, 1.83 |

\# Multi-plexed on one flow cell.

**Table 3.** Results for MinION sequencing.

| Species | Strain | # MinION Sequences | Base Pairs | Mean Length | Max Length | Genome Size | Mean Depth |
| --- | --- | --- | --- | --- | --- | --- | --- |
| *F. nucleatum* | 23726 | 13,904 | 81,303,330 | 5,847.5 | 55,886 | 2,299,539 | 35.4 X |
| *F. nucleatum* | 25586 | 25,240 | 111,179,924 | 4,404.9 | 84,482 | 2,180,101 | 60.0 X |
| *F. varium* | 27725 | 31,066 | 89,376,121 | 2,877.0 | 62,500 | 3,303,644<br>Plasmid:<br>42,814 | 27.0 X |
| *F. ulcerans* | 49185 | 20,313 | 42,439,006 | 2,089.3 | 74,841 | 3,537,675 | 12.0 X |
| *F. mortiferum* | 9817 | 37,658 | 118,090,402 | 3,135.9 | 83,201 | 2,716,766 | 43.5 X |
| *F. gonidiaformans* | 25563 | 58,596 | 105,220,409 | 1,784.7 | 87,683 | 1,678,881 | 62.7 X |
| *F. periodonticum* | 2_1_31 | 52,316 | 178,049,514 | 3,403.3 | 68,386 | 2,541,084 | 70.0 X |
| *F. necrophorum* | 1_1_36S | 66,335 | 129,875,928 | 1957.9 | 81,384 | 2,286,018 | 56.8 X |

### Genome assembly

Genome assemblies were carried out using the open-source software Unicycler^9^, resulting in complete chromosomes for each of the eight sequenced genomes. While both the Illumina and MinION sequencing runs produced far more data than necessary, data sets were split to utilize ample yet reasonable mean depth of coverage for 1.6 mb to 3.5 mb genomes. Using the mean depths of coverage for each genome described in Table 1 and Table 2, each genome can be constructed in 2-3 hours using a standard Macbook Pro laptop (2.8 GHz Intel Core i7). The utility of Unicycler therefore opens up a robust method for researchers without the need for a super computer to handle data processing. The details of all final assemblies are shown in Figure 2, and the public availability of the data at NCBI is detailed in Table 4. For consistent starts to the circular chromosome, each genome was rotated to have Gene 1, which encodes for the rod-shape determining protein MreC, in the reverse orientation as is seen for the beginning of the *F. nucleatum* subsp. *nucleatum* ATCC 25586 reference genome^8^.

**Figure 2:**
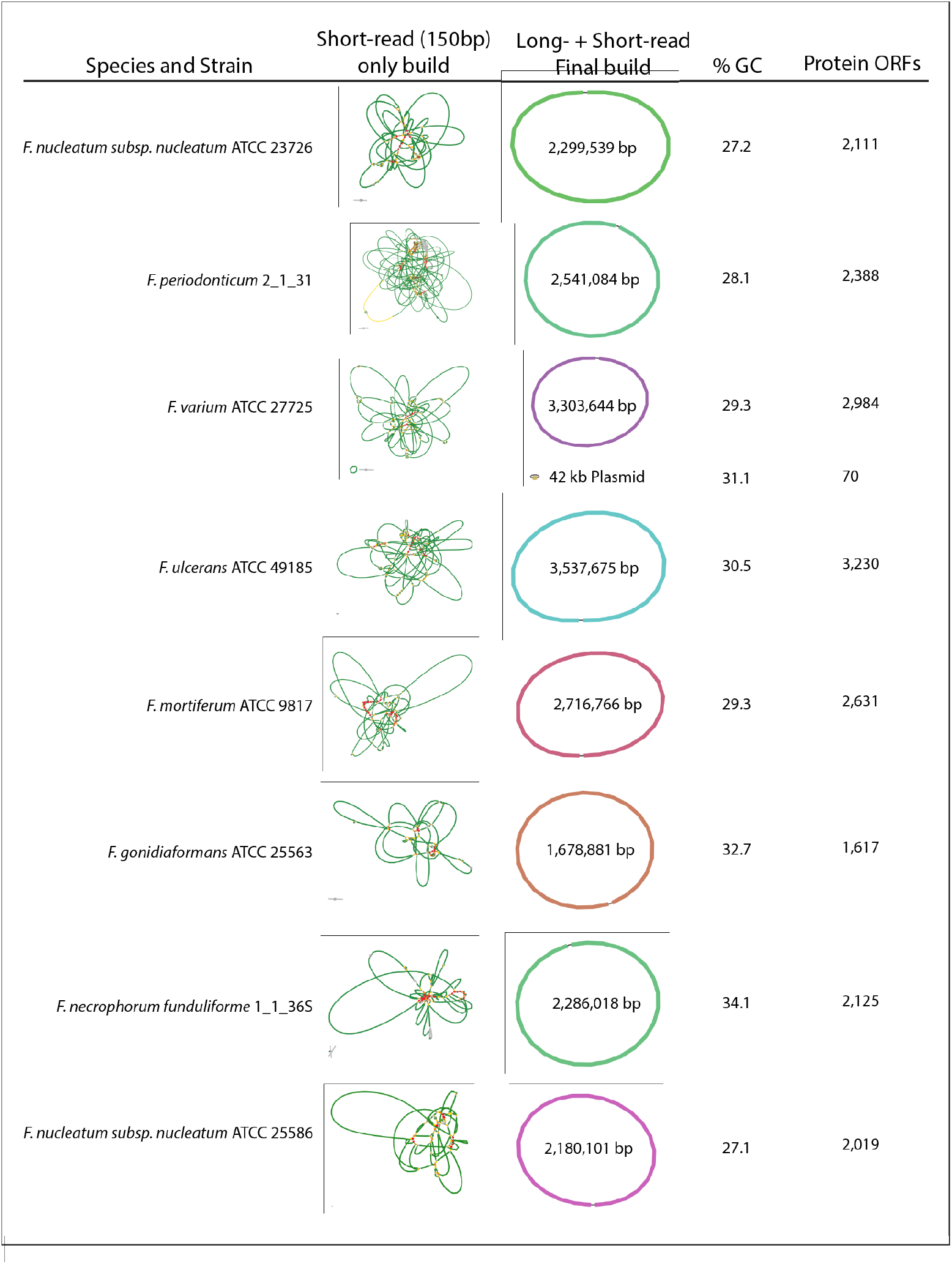
Genome assembly and Annotation of eight *Fusobacterium* genomes from seven species. Short-read only and complete genome assembly representations were created using Bandage^11^.

**Table 4.**
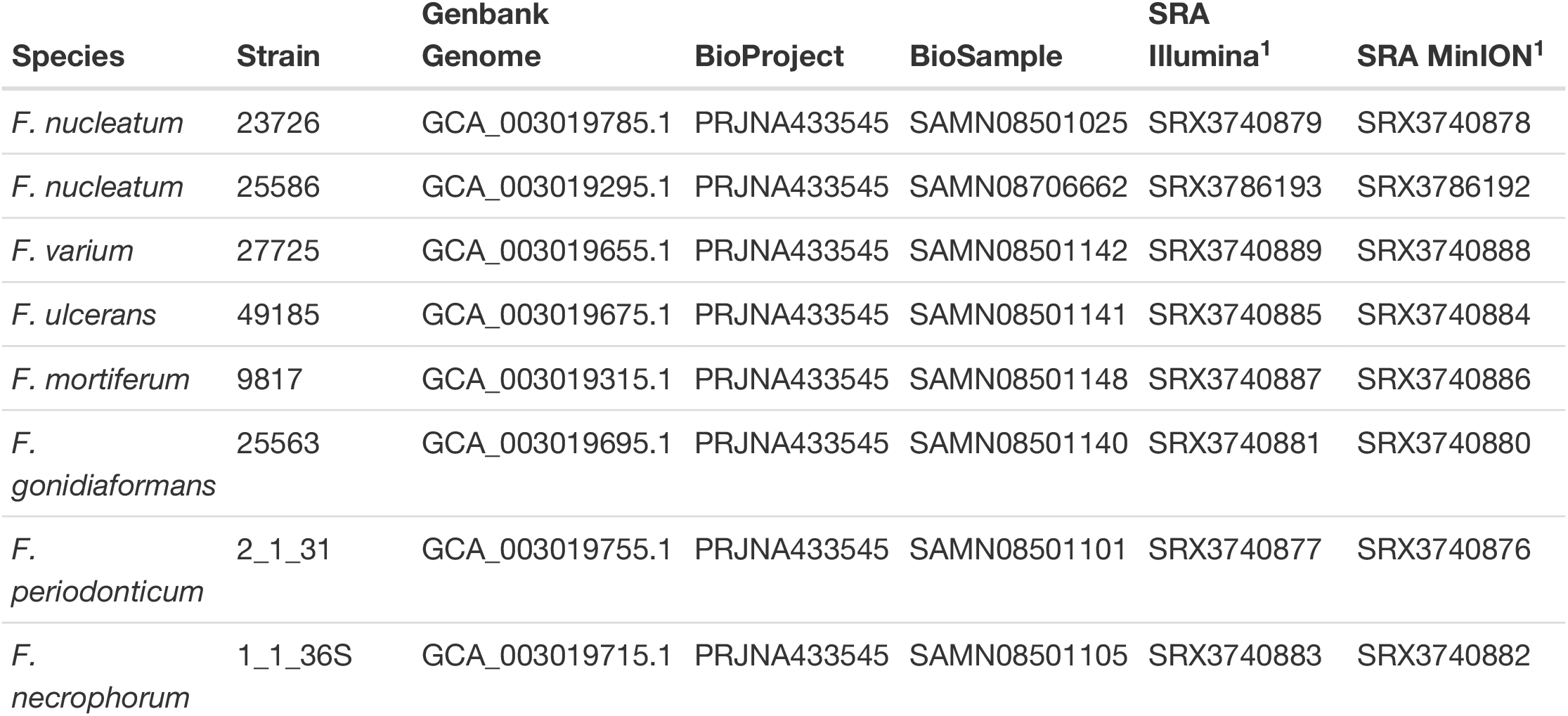
Data deposited at NCBI for all sequenced *Fusobacterium* strains.

^1^ Sequence Read Archive at NCBI

In addition, we believe that we have identified a previously undocumented 42 kb plasmid in *F. varium* ATCC 27725. To show the effectiveness of our genome assembly pipeline, Figure 3a shows the alignment of 67 contigs from the previous *F. nucleatum* subsp. *nucleatum* ATCC 23726 draft genome on our completed circular genome. We show that all contigs map, with our genome completing previous gaps. The accuracy of our genome when compared to mapped base pairs from the draft genome assembly at NCBI shows 99.99% base identification as determined by Geneious version 9.1.4. Strikingly, upon Geneious alignment of our *F. nucleatum* subsp. *nucleatum* ATCC 25586 genome with the previously complete genome deposited at NCBI (GCA_000007325.1), we discovered a ~ 452 kb genomic inversion (Figure 3b). This region is flanked on both ends by ~ 8 kb repeats that are likely the reason for the previous inability to discover this genomic feature. To validate this inversion, we aligned eight MinION reads (30-68 kb) that spanned this region, and show that these sequences confirm this genomic correction.

**Figure 3:**
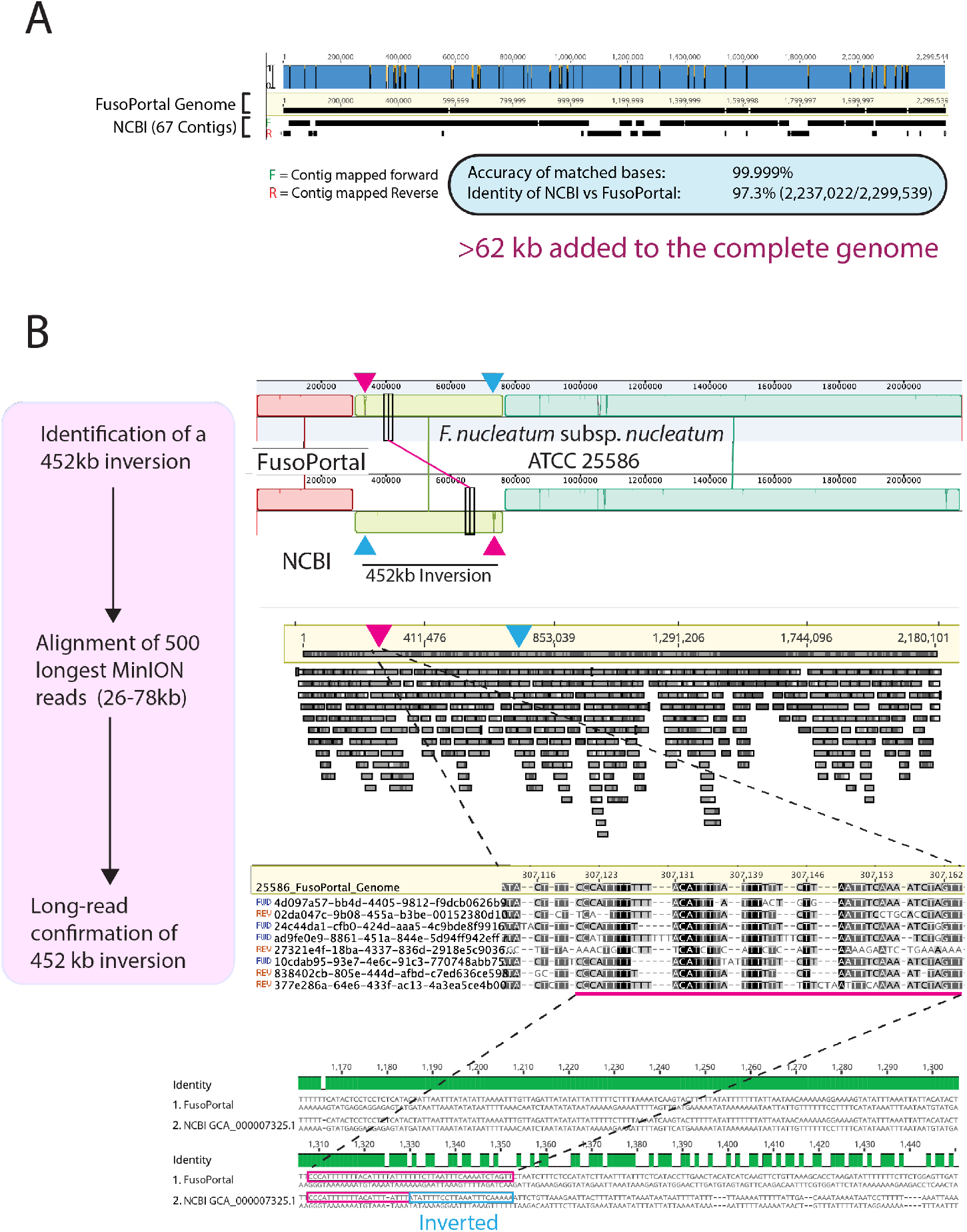
Analysis of *Fusobacterium nucleatum*. (A) Alignment of the complete *F. nucleatum* subps. *nucleatum* ATCC 23726 genome with the 67 contig draft assembly (Genbank: ADVK01000000). (B) Confirmation of a 452 kb genomeic inversion in the previous *F. nucleatum* subps. *nucleatum* ATCC 25586 genome assembly (Genbank: GCA_000007325.1).

### Open reading frame predictions

Gene predictions for protein encoding open reading frames were carried out using the bacterial specific program Prodigal via command line on a Mac^12^. Genes for tRNA encoding were predicted with Prokka^13^ using the KBase server^14^. rRNA were identified using Barrnap (Bacterial ribosomal RNA predictor). In addition, we used the CRISPRone web server to identify all CRISPR associated proteins and arrays, which consist of spacer and repeat regions. Details of each of these components are found on the FusoPortal repository. In each genome, protein encoding gene predictions by Prodigal and Prokka were in nearly complete agreement (data not reported). In addition, genome annotation for each genome was requested at NCBI upon data deposition into Genbank (Table 4).

### Software and code availability

All software and scripts used in this study have been described and properly referenced in previous methods sections.

## Data Records

Raw data and completed genomes for each of the eight *Fusobacterium* strains have been deposited at NCBI under the BioProject, BioSamples, Sequence Read Archives (SRA), and Genbank accession numbers detailed in Table 4.

## Technical Validation

CheckM^15^ on the Kbase^14^ server was used to check the quality of each genome using the reduced tree data set setting. An example of this data analysis is shown for strain *F. gonidiaformans* ATCC 25563 is shown in figure Figure 4, and data analysis for all genomes are available on the FusoPortal repository.

**Figure 4:**
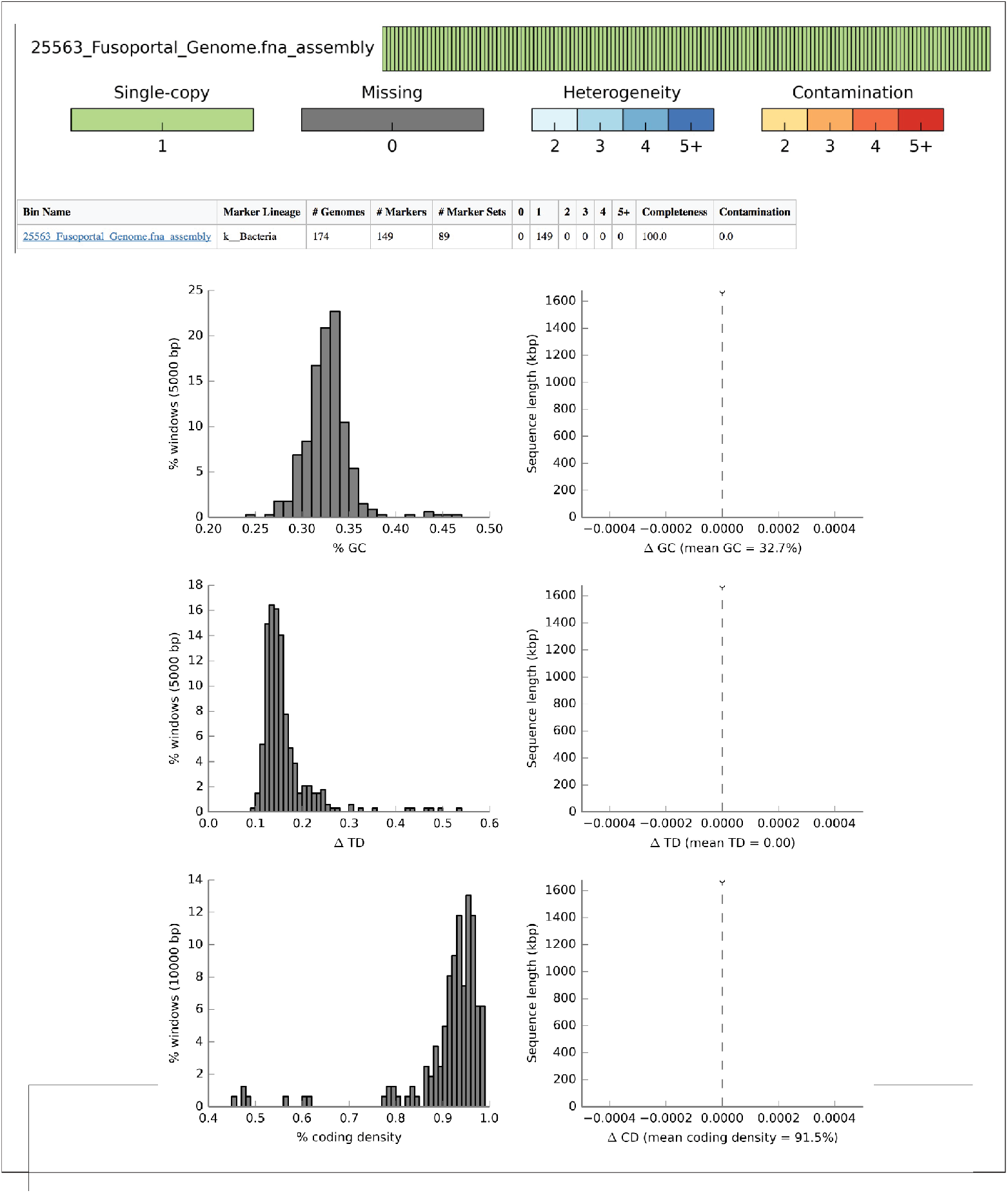
CheckM genome analysis of *F. gonidiaformans* ATCC 25563. CheckM analysis on the Kbase server shows that the *F. gonidiaformans* ATCC 25563 is of high quality and contains all core bacterial genes tested. Data analysis by CheckM for all eight *Fusobacterium* genomes described in this study are detailed on the FusoPortal repository.

## Usage Notes

The raw data, genome assemblies, and annotations can be accessed via the NCBI BioProject PRJNA433545, and further details of these files can be found in Table 4. In addition, all of this data is easily accessible in our newly implemented FusoPortal data repository or on our Open Science Framework database.

## Data Citations

1. *NCBI BioProject PRJNA433545* http://www.ncbi.nlm.nih.gov/bioproject/433545
2. *FusoPortal* http://fusoportal.org
3. *Open Science Framework* http://osf.io/2c8pv

## Acknowledgements

We would like to thank the laboratory of Emma Allen-Vercoe (University of Guelph) for providing many of the strains sequenced in this study. This work is supported by the USDA National Institute of Food and Agriculture.

## Author Contributions

S.M.T. performed all MinION sequencing, and wrote and edited the manuscript. K.K.L. prepared raw MinION sequences for genome assembly and wrote and edited the manuscript. R.E.S. assembled genomes and edited the manuscript. D.J.S. conceived and designed the experiments, assembled genomes, analyzed the data, and wrote and edited the manuscript.

## Competing Interests

The authors declare no competing financial interests.

## References

1. Dahya, V., Patel, J., Wheeler, M. & Ketsela, G. Fusobacterium nucleatum endocarditis presenting as liver and brain abscesses in an immunocompetent patient. Am. J. Med. Sci. 349, 284–285 (2015).

2. Signat, B., Roques, C., Poulet, P. & Duffaut, D. Fusobacterium nucleatum in periodontal health and disease. Curr. Issues Mol. Biol. 13, 25–36 (2011).

3. Castellarin M. et al. Fusobacterium nucleatum infection is prevalent in human colorectal carcinoma. Genome Res. 22, 299–306 (2012).

4. Kostic A. D. et al. Genomic analysis identifies association of fusobacterium with colorectal carcinoma. Genome Res. 22, 292–298 (2012).

5. Kostic A. D. et al. Fusobacterium nucleatum potentiates intestinal tumorigenesis and modulates the tumor-immune microenvironment. Cell Host Microbe 14, 207–215 (2013).

6. Rubinstein M. R. et al. Fusobacterium nucleatum promotes colorectal carcinogenesis by modulating E-cadherin/ β-catenin signaling via its FadA adhesin. Cell Host Microbe 14, 195–206 (2013).

7. Yu T. et al. Fusobacterium nucleatum promotes chemoresistance to colorectal cancer by modulating autophagy. Cell 170, 548–563.e16 (2017).

8. Kapatral V. et al. Genome sequence and analysis of the oral bacterium fusobacterium nucleatum strain ATCC 25586. J. Bacteriol. 184, 2005–2018 (2002).

9. Wick, R. R., Judd, L. M., Gorrie, C. L. & Holt, K. E. Unicycler: Resolving bacterial genome assemblies from short and long sequencing reads. PLoS Comput. Biol. 13, e1005595 (2017).

10. Wick, R. R., Judd, L. M., Gorrie, C. L. & Holt, K. E. Completing bacterial genome assemblies with multiplex MinION sequencing. Microb Genom 3, e000132 (2017).

11. Wick, R. R., Schultz, M. B., Zobel, J. & Holt, K. E. Bandage: Interactive visualization of de novo genome assemblies. Bioinformatics 31, 3350–3352 (2015).

12. Hyatt D. et al. Prodigal: Prokaryotic gene recognition and translation initiation site identification. BMC Bioinformatics 11, 119 (2010).

13. Seemann T. Prokka: Rapid prokaryotic genome annotation. Bioinformatics 30, 2068–2069 (2014).

14. Arkin A. P. et al. The DOE systems biology knowledgebase (KBase). bioRxiv 096354 (2016).

15. Parks, D. H., Imelfort, M., Skennerton, C. T., Hugenholtz, P. & Tyson, G. W. CheckM: Assessing the quality of microbial genomes recovered from isolates, single cells, and metagenomes. Genome Res. (2015).

